# Histological morphometric correlates of MRI-defined cortical thinning in temporal lobe epilepsy

**DOI:** 10.64898/2026.10.22.761130

**Authors:** Joanna Bartkiewicz, Fenglai Xiao, Yau Mun Lim, Maritchka Ryniejska, Hanaa El Hachami, Alicja Mrzyglod, Joan Liu, Anna Miserocchi, Andrew W McEnvoy, John S Duncan, Matthias Koepp, Maria Thom

## Abstract

Large-scale MRI studies have consistently demonstrated progressive and age-accelerated neocortical thinning in temporal lobe epilepsy (TLE). These findings are frequently interpreted as evidence of cumulative neurodegeneration. However, the histopathological substrates underlying MRI-confirmed cortical thinning in epilepsy remain poorly defined. In particular, it is unknown whether cortical thinning reflects irreversible neuronal loss or potentially reversible epilepsy-related network alterations in cortical microstructure.

We analysed the resected middle temporal gyrus from 117 individuals undergoing surgery for temporal lobe epilepsy, including cases with (n= 74) or without hippocampal sclerosis (HS) and 12 cases with additional temporal lobe sclerosis (TLS, or Focal Cortical Dysplasia IIIa), defined by a distinct pattern of cytoarchitectural reorganisation and gliosis involving superficial layers. All cases were stratified by the presence or absence of pre-operative MRI-defined cortical thinning of the middle temporal gyrus, quantified using CAT12 relative to healthy controls. On NeuN-immuno-labelled sections using automated image analysis we assessed neuronal spatial distributions relative to their size and packing density; relative cortical layer thickness and sulcal-gyral cytoarchitectural variation were also quantified.

MRI-defined cortical thinning was not associated with diffuse neuronal loss. Instead, thinning correlated with increased neuronal density and altered cell-size spatial organisation in superficial and deep layers, particularly in right temporal lobe epilepsy cases with HS. There was limited evidence of layer-specific laminar thinning, being confined to layer VI in the right TLE/HS group only. Of note, TLS cases did not show either cortical thinning on MRI or laminar thinning on histology compared to cases without TLS but altered neuronal densities and composition that was not confined to the superficial cortex. MRI cortical thinning was also characterised by attenuation of normal sulcal/gyral cytoarchitectural variations suggesting disruption of cortical architectural gradients. In contrast, cases with HS but without MRI cortical thinning demonstrated loss of the expected inter-hemispheric cytoarchitectural asymmetry, indicating a distinct pathological process dissociated from thinning. Clinical-pathological correlations were lateralised, including age of onset, further supporting distinct mechanisms contributing to MRI cortical thinning between left and right TLE.

These findings challenge the prevailing assumption that MRI-defined cortical thinning in epilepsy reflects progressive neuronal degeneration. Rather than laminar neuronal loss and atrophy, thinning was associated with altered neuronal packing and cell-size distributions between layers. This points towards altered neuropil volumes, including myelin content and dendritic remodelling, in turn suggesting potentially reversible dynamic seizure- or network-related processes.

## Introduction

MRI-defined measures of regional cortical thickness are widely used to characterise and monitor disease progression in neurodegenerative disease^1,2^, psychiatric conditions^3,4^ and normal aging^5^. In temporal lobe epilepsy (TLE), similar approaches have consistently demonstrated progressive and age-accelerated neocortical thinning extending beyond the seizure focus. These findings support the concept of TLE as a disturbed network disorder^6^ and, in some patients, a progressive process^7^ associated with surgical outcome and cognitive decline in later life^8^. However, a fundamental, unresolved question remains: does MRI-defined cortical thinning in epilepsy represent irreversible neuronal loss or does it reflect cumulative seizure-related and network-driven remodelling of cortical structure.

With regard to the temporal neocortex in TLE, early voxel-based morphometry (VBM) studies identified grey matter volume reduction, more marked ipsilateral to the seizure focus and spanning superior to inferior gyri^9^. Subsequent thickness analyses reported reductions of 5 to 15%, again predominately ipsilateral^10^, noting differences between cases with or without hippocampal sclerosis (HS)^11^ and divergence from normal aging trajectories^12,13^. These findings were interpreted as evidence of epileptogenic networks involvements and cumulative seizure-related atrophy. Advances in computational neuroimaging, including FreeSurfer and the Computational Anatomy Toolbox (CAT12), have enabled large multicentre analyses. Studies from the ENIGMA (Enhancing NeuroImaging Genetics through Meta-Analysis) Epilepsy consortium confirmed ipsilateral temporal lobe thinning in left but not right TLE associated with HS (TLE/HS)^6^. Subsequent studies identified bilateral superior temporal cortical atrophy related to highly connected network hubs^14^, with more pronounced ipsilateral effects, particularly in right TLE^15^. Both cross-sectional and longitudinal MRI studies further supported accelerated age-related grey matter loss in epilepsy, particularly in people over the age of 55^16^, at rates exceeding normal aging^7,13^.

At the same time, accumulating evidence suggests that MRI-defined cortical thickness may not exclusively index irreversible neuronal degeneration. Successful temporal lobe surgery has been associated with attenuation of progressive atrophy^17^ and, in some cases, regional cortical thickening post-operatively^18^. Increases in grey matter volume or cortical thickness have also been reported pre-operatively in TLE^19^, following generalised seizures^20^ after electroconvulsive therapy^21^ and in association with cardiovascular fitness^22^. Together these observations suggest that cortical thickness measures may, under certain conditions, reflect activity-related dynamic and potentially reversible microstructural changes rather than fixed neuronal loss. The histopathological substrates of MRI-defined cortical thinning in TLE remain poorly defined.

Common neocortical abnormalities in TLE/HS surgical specimens include subpial and intracortical gliosis^23^, oligodendrogliosis^24^, neuronal hypertrophy^25,26^, microvasculature alterations^24^ and cortical myeloarchitectural changes^26^ several of which may influence MRI-defined contrast and thickness measurements and may not necessarily represent irreversible degeneration^27,28^. Temporal lobe sclerosis (TLS), present in 11% of TLE/HS cases, represents a distinct cytoarchitectural pattern characterised by neuronal reduction in layers II/III with clustering and hypertrophy of neurones in upper layer II and associated myeloarchitectural abnormalities^26^. Due this pattern of disorganisation, TLS is classified as one subtype of focal cortical dysplasia (FCD) type IIIA in the ILAE classification^29^.

Direct MRI-histology studies in TLE remain scarce and mainly based on small case numbers. VBM analysis in 13 TLS cases did not detect significant grey matter reductions compared with controls^30^, MRI-histology co-registration studies suggested inverse relationships between T2 and NeuN labelling indices in the middle temporal gyrus (MTG), interpreted as reflecting altered neuropil volume or cortical myelin fraction^27^, while correlations between MRI cortical thickness and histological layer II/III thickness have been reported in 36 cases^31^. More recent work implicates microglial activation as a modifying factor in cortical thinning^32^. Nonetheless, whether MRI-defined cortical thinning in TLE reflects neuronal depletion, neuropil contraction, altered laminar organisation, or a combination of these processes remains uncertain.

In this study we aimed to examine the histological correlates of MRI-defined cortical thinning in a large, well-characterised surgical cohort of TLE. By combining automated MRI-defined cortical thickness measures with quantitative NeuN-based cytoarchitectural analysis of MTG, we sought to determine whether cortical thinning primarily reflects neuronal loss or altered cortical organisation, and to explore potential lateralised mechanisms underlying these changes.

## Materials and methods

### Case Selection

We included 117 TLE cases from the Epilepsy Brain and Tissue Bank (ESBTB) at University College London that had undergone standard anterior temporal lobe resection and hippocampectomy between 2003 and 2023 at the National Hospital for Neurology and Neurosurgery (London, UK). As part of presurgical investigations, all cases underwent structural MRI on 3T GE Discovery MR750 Scanner using a 3D T1-weighted magnetisation prepared rapid acquisition gradient echo (MPRAGE) sequence with TE/TR/TI: 3.1/7.4/400 ms, voxel size: 1.0×1.0×1.0 mm. Cases were selected based on both optimal neuroimaging and availability of resected tissue for research with the ESBTB through its ethical approval (NRES 23/SC/0002).

The pathology diagnosis in the selected 117 cases included 73 individuals with left or right mesial TLE with confirmed HS Type I pathology (mean age of 39.12 years ±10.54; 29 males; Supplementary Table 1), 6 with HS Type II, 2 with HS Type III, 12 with temporal lobe sclerosis (TLS, also termed as FCD IIIa ^33^) and 24 with no lesional pathology (including no definitive HS/ hippocampal gliosis only). Patients with dual pathology,

e.g. tumours, cavernomas or FCD type II were excluded. The demographic data and seizure history of the cohort, including duration of epilepsy, laterality and seizure types, are summarised in Table 1 and Supplementary Table S2 (additional case detail in Supplemental Table S1).

**Table 1.** Clinical Details for CT groups.

|  | **Left**<br>**TLE**<br>**HCT** | **Left**<br>**TLE**<br>**LCT** | **Right**<br>**TLE**<br>**HCT** | **Right**<br>**TLE LCT** |
| --- | --- | --- | --- | --- |
| **Cases, n** | 11 | 29 | 11 | 26 |
| **Age at Surgery** | 41.64<br>(11.27) | 37.00<br>(9.94) | 43.55<br>(14.41) | 38.54<br>(8.74) |
| **Age of Seizure Onset** | 8.41<br>(7.68) | 10.95<br>(9.84) | 18.64<br>(14.08) | 12.04<br>(7.49) |
| **Range Age of Seizure Onset** | 0.25-26 | 0.5-34 | 2-50 | 1-30 |
| **Duration of Epilepsy** | 33.23<br>(14.03) | 26.05<br>(12.28) | 24.91<br>(14.43) | 26.50<br>(11.31) |
| **Average Frequency of Seizures during 12 months preop** | 12.36 | 13.54 | 8.00 | 5.55 |

Abbreviations: SD = Standard deviation; TLE = temporal lobe epilepsy; HCT = high cortical thinning; LCT = low cortical thinning Age at Surgery, Age at Seizure Onset and Duration of epilepsy are presented as average in years (SD).

For the MRI analysis only, epilepsy cases were compared to a group of 70 healthy controls (HC) (44 female) with no history of neurological or psychiatric disease [mean age of 39.77± 12.82 for females, 41.63 ± 12.91 for males).

### MRI Processing and Analysis

All images were corrected for movement and distortion artifacts using Computational Anatomical Tool (CAT12^34^) on SPM12 and then spatially normalised using CAT12 to MNI space. Segmentation and parcellation of T1 was performed using Desikan-Killiany atlas. Using CAT12, individuals with epilepsy were analysed against HC to measure the cortical thickness for each parcellated region of the brain, including MTG. Patients were grouped according to MRI-based cortical thickness relative to healthy controls in MTG region, with the cutoff set at (mean – 1SD) (for left and right TLE: 2.67 and 2.73mm, respectively). Those with lower thickness were classified as high cortical thinning (HCT), and those above as low cortical thinning (LCT; Supplementary Table S1). Due to anatomical differences between the hemispheres, we grouped left (L) and right (R) TLE patients separately, thus forming four groups: LHCT (n=16), LLCT (n=40), RHCT (n=18) and RLCT (n=43), collectively known as “CT Groups” (Fig. 1I-L). The aim was to obtain equal relative representation of LCT an HCT in both left and right TLE in view of differences in left/right functional connectivity^35^, clinical epilepsy presentations^36^ and cortical thicknesses in MRI epilepsy studies^6^. The time interval between the MRI and the surgical resection was also recorded (Supplementary Table S2).

**Figure 1.**
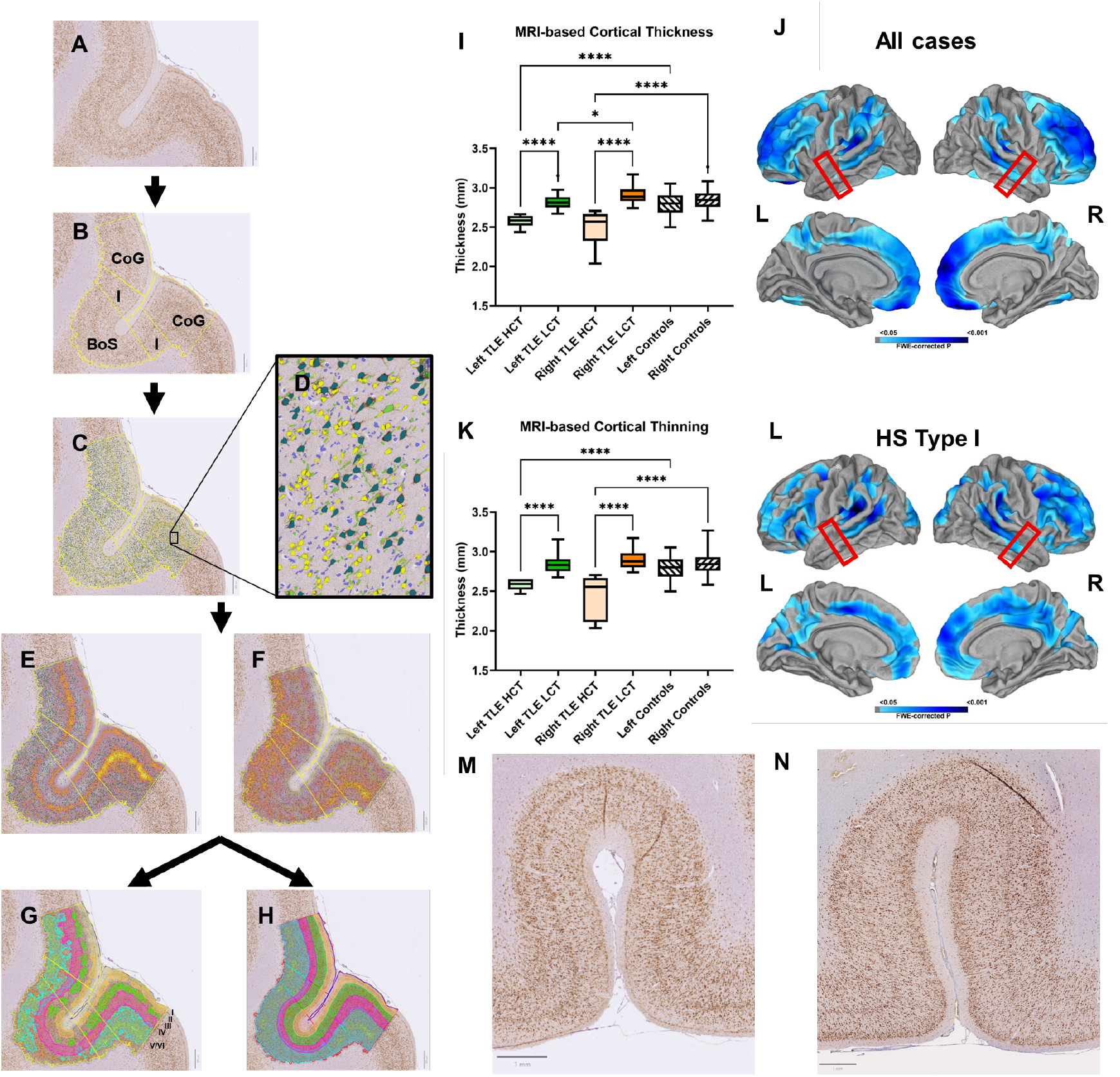
Cortical thinning across scales: MRI-based cortical thinning and NeuN^+^ IHC staining. **A-H** show a flow diagram of the pipeline for annotating digital slides, where (**A**) Brightfield digitalised slides are loaded into QuPath software. (**B**) Regions of interest (ROIs) are annotated; CoG - representing the crown of gyrus on either side of the sulcus, I - the intermediate region including the mid-way region between crown of gyrus and sulcal depth and BoS - the bottom of sulcus. (**C**) NeuN^+^ cells were detected in QuPath based on cell area: small S-NeuN^+^ (yellow) between 50–149 µm^2^, and large L-NeuN^+^ (dark green) between 150–300 µm^2^ with cells outside these parameters excluded (bright green), and negative cells (blue). (**D**) shows magnification of rectangle in (**C**). After detecting and labelling NeuN^+^ cells, we generated density heat maps for **(E)** S-NeuN^+^ and (**F**) L-NeuN^+^. These were then used to outline the cortical layers, either in a (**G**) automated version, where QuPath outlines the edges of the heat map, or (**H**) manual version, where the density heat maps were guiding the annotation, but also followed the natural shape of the laminar layers. **(I)** shows one-way ANOVA analysis comparing MRI-based cortical thickness of the entire cohort (n=117) divided into appropriate CT groups and control groups. (**J**) shows one sample t-test comparison of all patients with epilepsy against a group of controls. (**K**) one-way ANOVA analysis comparing MRI-based cortical thickness of patients with HS type I pathology only divided into appropriate CT groups and control groups. (**L**) shows one sample t-test comparison of all patients with HS type I pathology against a group of controls. Challenge of identifying cortical thinning in histological tissue can be seen in (**M-N**), where it shows examples of two NeuN^+^ brightfield photomicrographs from (**M**) HCT and (**N**) LCT groups. Red boxes in **J** and **L** indicate the location of temporal lobe analysed immunohistochemically. All tests were corrected for multiple comparisons, significance is shown with asterisk, where * *p*<0.05 and **** *p*<0.0001. HCT = high cortical thinning; LCT = low cortical thinning; HS = hippocampal sclerosis.

### Immunohistochemistry

Temporal lobes were all handled and processed using a standardised protocol as previously described^37^. In this study, the coronal block 1 to 1.5 cm caudal to the temporal pole was selected from each case, which shows the full representation of all gyri with minimal surgical and edge handling artefacts. Formalin-fixed paraffin-embedded sections were cut at 14µm and immunostaining carried out with Discovery Ultra autostainer (Ventana, Roche Roche Diagnostic Limited, UK), sections were incubated for an hour with neuronal marker NeuN at room temperature and amplified using Amplification kit (05266114001). Slides

### Image Analysis

Scans of NeuN-immunolabelled sections were imported into QuPath v0.6.0 (open source software for bioimage analysis^38^) and the MTG and the adjoining superior or inferior temporal sulcus of each case were analysed (see Supplementary Table S4). The MTG of each case was subdivided manually into three regions of interests (ROIs) according to the curvature of the gyrus to include the crown, the lateral intermediate area, and the bottom of the sulcus. Each ROI consisted of the full thickness of the cortex, delineated by the pial surface and white matter boundary at the superior and inferior aspects respectively (see Fig. 1A-B). Tissue artefacts, such as tears and folds, were excluded from ROI.

All NeuN immunopositive cells with an intensity threshold of 0.1 were quantified using the Positive Cell Detection module on QuPath, followed by further classification according to cell size (small or S-NeuN+, 50-149 µm2; large or L-NeuN+, 150-300 µm2) to utilise the differential neuronal domination between laminae (see Fig. 1C-D). A script was written to generate heat maps based on relative densities of S-NeuN+ and L-NeuN+ (Fig. 1E-F). When superimposed onto the ROIs, the heat maps identified and outlined four cortical laminae (superior cortical layers II and III, and deep cortical layers, IV and V/VI; Fig. 1G-H). The density maps layer V and VI were not separable and these hereafter are termed layer VI. The borders of the cortical layers were subsequently defined by both a semi-manual (Fig. 1H) as well as a fully automated (Fig. 1G) method, the later which extrapolated or imputed regions or gaps in the density map due to immunostaining artefacts, less intense labelling or regions of low cell density, e.g. layer II in TLS cases. Layers II and III were also grouped were then incubated for 24min in a solution containing anti-mouse HRP secondary antibody (OmniMap 05269652001) followed by chromogen detection using 3,3’-diaminobenzidine 28 (DAB) (ChromoMap DAB, 05266645001, Roche), then counterstained with haematoxylin (Haematoxylin II, 05277965001). Sections were then scanned at 40x with digital slide scanner (Hamamatsu NanoZoomer S360, Hamamatsu Photonics). as superficial cortical layers and layers IV and V/VI, deep cortical layers.

Due to the inherent curvature of the human cortex and non-orthogonal or tangential tissue sections, absolute measurements of cortical and laminar thickness based on histology sections is not possible. Therefore, we used relative metrics to enable valid cross-case comparisons. For each layer, we computed: (1) Percentage layer thickness (layer thickness): distance between the 10th and 90th percentile of all NeuN^+^ cells in that layer to the defined pial border divided by the total ROI thickness (pia to white matter), (2) percentage S-NeuN^+^ and L-NeuN^+^ based on the proportion of S-NeuN^+^ or L-NeuN^+^ within a layer relative to all detected NeuN^+^ cells of the same size range within that ROI and (3) cell density metrics for S-NeuN^+^ (density S-NeuN^+^) and L-NeuN^+^ (density L-NeuN^+^), defined as the number of detected NeuN^+^ cells per layer area (µm^2^).

### Statistical Analysis

Statistical analysis was conducted with GraphPad Prism (Prism 10, Version 10.4.2) and R (RStudio 2024.04.2). Associations between epilepsy-related clinical variables (age, sex, duration, history of previous injury, seizure type, monthly frequency of seizures and post-operative outcome) and histopathological measures were examined (Table 1 and Supplementary Table S2).

For all dependent variables, descriptive analysis and distributions were assessed for normality using visual inspection and formal tests. Based on these assessments, either parametric or non-parametric statistical tests were applied as appropriate.

Where multiple comparisons were performed, p-values were adjusted using Bonferroni correction for parametric analyses or Dunn’s correction following non-parametric tests. Adjusted: *p<*0.05 were considered significant. Statistical analyses were conducted in consultation with the UCL statistical support service.

### Confounder Analysis

To identify potential confounding relationships between clinical variables, pairwise comparisons were performed across all clinical factors using non-parametric tests appropriate to variable type. Spearman’s rank correlation was used to assess associations between continuous variables, Mann–Whitney U or Kruskal–Wallis tests were used for comparisons between continuous and categorical variables, and Chi-square tests were applied for associations between categorical variables.

Variables showing significant (*p* < 0.05) and strong (r > 0.6) correlation coefficients were considered potential confounders and included as covariates in subsequent regression analyses.

### Group comparison Analysis

For proportional dependent variables, relative cortical layer thickness, percentage of small NeuN-positive neurones (S-NeuN^+^), and percentage of large NeuN-positive neurones (L-NeuN^+^), representative values were calculated for each patient as weighted averages across all regions of interest (ROIs), accounting for ROI size.

Between-group comparisons were performed using one-way ANOVA or Kruskal–Wallis tests, depending on normality. Interactions between cortical thinning group and cortical layer or ROI were assessed using repeated-measures two-way ANOVA. Agreement between automated and manual layer outlining was evaluated using the Bland–Altman method.

### Regression Analysis

For neuronal density measures, linear regression was applied to S-NeuN^+^ density following confirmation of

### Quantitative pathology findings

#### Comparison and correlation of clinical factors

normality, while L-NeuN^+^ density was analysed using gamma regression with a log link function to accommodate positively skewed distributions. One subgroup (right hemisphere cortical thinning) showed mild left-skewness across layers; however, this did not violate model assumptions, which were verified using diagnostic plots and residual analyses.

Proportion-based outcomes (relative layer thickness, percentage of S-NeuN^+^, and percentage of L-NeuN^+^) were analysed using beta regression with a logit link, implemented using the betareg package in R. Predictor variables included MRI-defined cortical thickness, age at seizure onset, age at surgery, duration of epilepsy, monthly seizure frequency, post-operative outcome, sex, history of previous injury and seizure type. Categorical variables were coded using males, childhood convulsions and focal impaired awareness seizures as reference categories.

Only models demonstrating acceptable goodness-of-fit, assessed using Akaike and Bayesian Information Criteria (AIC and BIC) are reported in the Results section. Effect estimates are presented as regression coefficients (ß) with associated *p*-value, alongside R^2^ values for linear regression models and *pseudo*-R^2^ values for beta regression models (Supplementary Table S5).

## Results

### Qualitative histopathological observations

On NeuN-immunostained sections, cases with MRI-defined HCT or LCT could not be distinguished by visual inspection alone with all specimens demonstrating well-preserved cortical lamination (Fig. 1M and N). TLS patterns, characterised by neuronal loss in layer II and III (Fig. 2C), was identified in 12 cases and were evenly distributed between HCT and LCT groups (n=6 each) and occurred on both the left (n=8) and right (n=4) sides (Table 1).

**Figure 2.**
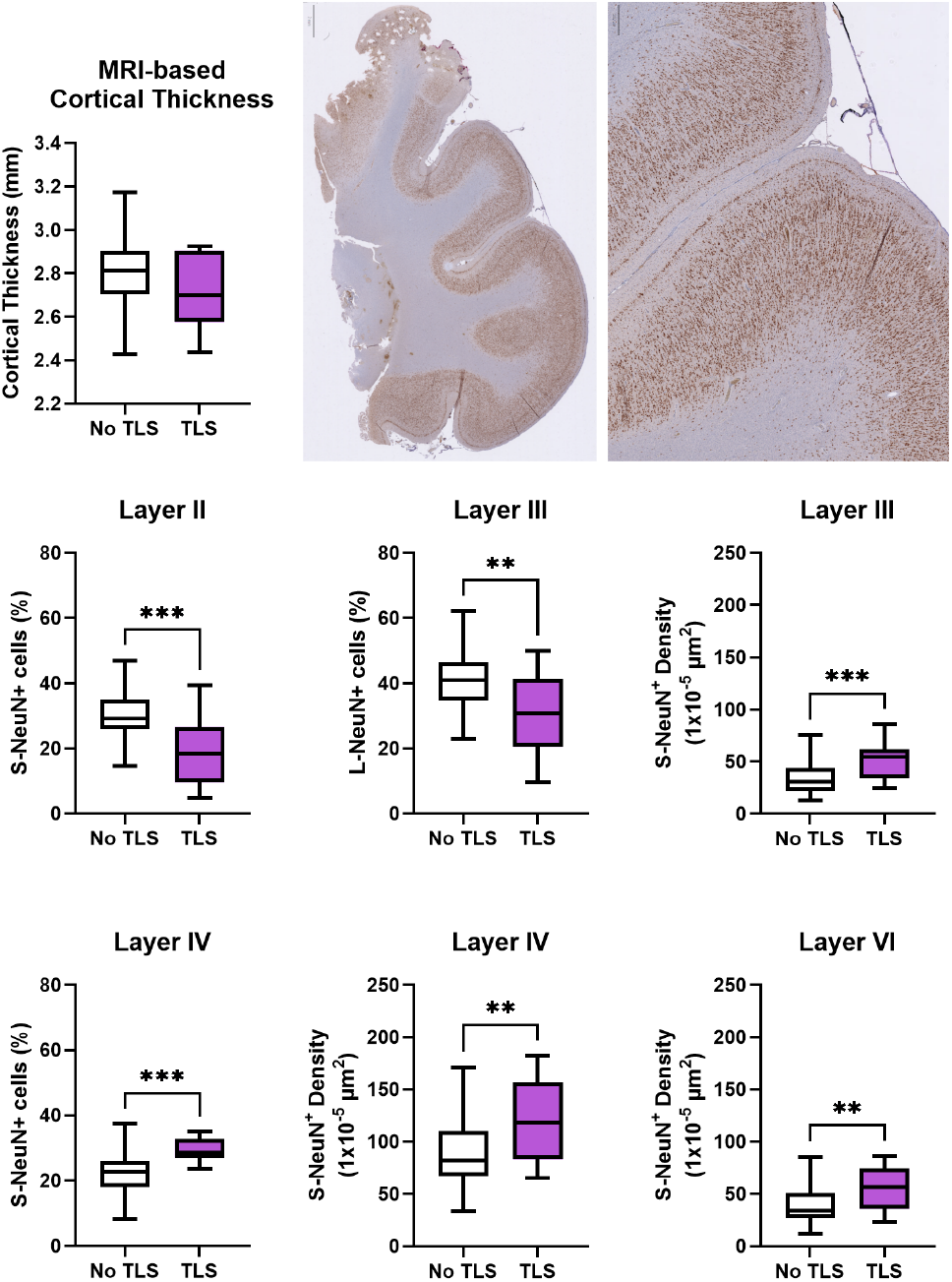
Cortical thinning temporal lobe sclerosis. Analysis of cases with TLS pathology in comparison to all other cases irrespective of left/right resection side. (A) shows no difference in MRI-based cortical thickness between groups. (B) shows brightfield photomicrograph of NeuN+-positive staining in an example of TLS pathology. (C) magnified (B) image to show to appreciate the characteristic thinning of Layer II resulting in a thin band. (D-I) shows Mann-Whitney test comparison analysis of TLS against no-TLS groups for dependent variables. Only significant results are shown here, for all comparisons, see Supplementary Fig. S6. (D-F) show significant results for superficial cortical layers, where (D) shows a significantly lower percentage of S-NeuN+ in TLS group in Layer II (p<0.0001), (E) significantly lower percentage of L-NeuN+ Layer III (p=0.003) and (F) significantly higher density of S-NeuN+ in Layer III (p=0.001). (G-I) show significant results in deep cortical layers, where in TLS cases (G) shows significantly higher percentage of S-NeuN+ in Layer IV (p<0.0001), (H) significantly higher density of S-NeuN+ in Layer IV (p=0.006) and (I) in Layer VI (p=0.003). All tests were corrected for multiple comparisons, significance is shown with asterisk, where * p<0.05, ** p<0.01, and *** p<0.001. TLS = temporal lobe sclerosis.

No significant differences in clinical variables were observed between cortical thickness groups or between pathology types, either in the full cohort (n=117) or when restricting analysis to HS type I cases (n=73; Supplementary Fig. S1, S2).

Correlation analysis using quantitative laminar data averaged across gyral ROIs revealed lateralized associations between clinical factors and cytoarchitectural measures. In left TLE, density of both small (S-NeuN^+^) and large (L-NeuN^+^) neurons were negatively correlated with duration of epilepsy in layer III (*p*=0.025, *p*=0.038, respectively), IV (*p*=0.047, *p*=0.034, respectively), and VI (*p*=0.031, *p*=0.009, respectively; Supplementary Fig. S3). Additional positive correlations were observed primarily in layers III and IV, including associations between sex with layer thickness (*p*=0.034), MRI-based cortical thickness and percentage S-NeuN^+^ (*p=*0.037), and monthly seizure frequency and percentage of L-NeuN^+^ (*p*=0.016).

In right TLE, significant correlations were fewer and predominantly involved the percentage of L-NeuN^+^ neurones, which was negatively correlated with age at surgery (*p*=0.043) and epilepsy duration (*p*=0.016; layer III), and positively correlated with a history of initial precipitating injury (*p*=0.036; layer II) (Supplementary Fig. S3).

When analysis was restricted to HS Type I cases, the pattern of lateralisation shifted. In right TLE with HS type I, duration of epilepsy showed strong negative correlations with percentage S-NeuN^+^ and L-NeuN^+^ neurons in layers III (*p*=0.039, *p*=0.016, respectively) and IV (*p*=0.012, *p*=0.03, respectively; Supplementary Fig. S4). MRI-defined cortical thickness was negatively correlated with the density of S-NeuN^+^ neurons in layers II (*p*=0.007), IV (*p*=0.005), and VI (*p*=0.044), and with L-NeuN^+^ density in layers IV (*p*=0.032) and VI (*p*=0.02). Full correlation results are provided in Supplementary Table S6.

#### Comparisons of HCT and LCT groups: whole cohort Relative laminar thickness

Relative cortical layer thickness did not differ between groups across any layers (Supplementary Fig. S5A–D).

#### L-NeuN^+^ distribution

While the proportion of L-NeuN^+^ did not differ between cortical thickness groups, significant differences were observed in neuronal density. Right-sided HCT (RHCT) cases showed higher L-NeuN^+^ density than left-sided (LHCT) in layers II and III (both *p*=0.01). RHCT also showed higher L-NeuN^+^ density than right-sided LCT (RLCT) in layers II (*p*=0.01) and IV (*p*=0.01; Supplementary Fig. S5M-O).

#### S-NeuN^+^ distribution

Proportional differences in S-NeuN^+^ were confined to superficial layers. LHCT showed a lower percentage of S-NeuN^+^ in layer II compared with left-sided LCT (LLCT; *p*=0.02) and a lower percentage in layer III compared with RHCT (*p*=0.007). Density differences were minimal, except in layer IV, where LHCT showed higher S-NeuN^+^ density than LLCT (*p*=0.01), while LLCT showed lower density than RLCT (*p*=0.01; Supplementary Fig. S5I-J, S).

#### Temporal lobe sclerosis group

TLS cases did not differ from non-TLS cases in MRI-based cortical thickness (Fig. 2A) or relative layer thickness (Supplementary Fig. S6A–D). However, marked differences in neuronal composition were observed. In superficial layers, TLS showed a lower proportion but higher density of S-NeuN^+^ (*p*<0.001 and *p*=0.001 respectively) alongside a reduced proportion of L-NeuN^+^ (*p*=0.003, Fig. 2D-F). In contrast, deeper layers showed the opposite pattern, with higher proportions and densities of S-NeuN^+^ neurones in TLS, particularly in lavers IV and VI (Fig. 2G-I).

#### Comparisons of HCT and LCT groups: HS type I

Given the distinct observations above with TLS, subsequent analysis was focused on cases with only HS type I (n=73).

#### Relative laminar thickness

Layer VI was significantly thicker in RLCT compared with RHCT (*p* = 0.02; Fig. 3).

**Figure 3.**
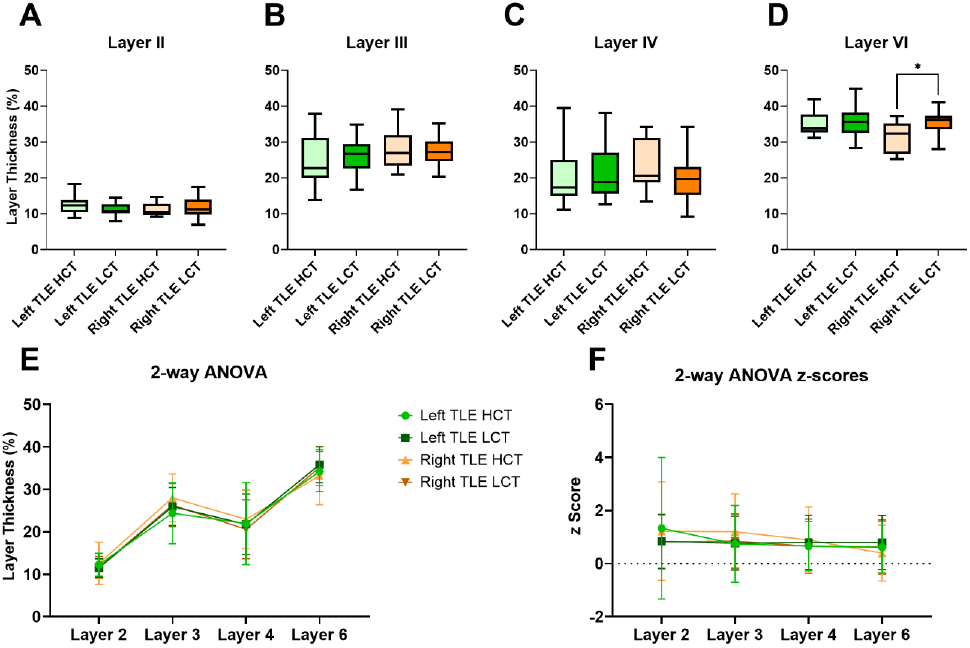
Comparison of layer thickness between CT Groups (TLE-HS Type I cases). A-D shows one-way ANOVA analysis between CT groups for each cortical layer, where comparisons are made between groups that had resection surgery on the same side and between same contralateral CT category, i.e. Left TLE HCT vs Right TLE HCT. (E) and (F) shows 2-way ANOVA analysis of repeat measures of Layer Thickness, accounting for multiple cortical layers measured within the same patient across CT groups (E) and z-scores (F). All tests were corrected for multiple comparisons, significance is shown with asterisk, where * p<0.05. HCT = high cortical thinning; LCT = low cortical thinning; HS = hippocampal sclerosis.

#### L-NeuN^+^ distribution

Proportional differences in L-NeuN^+^ neurones were modest, with RHCT showing higher proportion than RLCT in layer II (adjusted. *p* = 0.05), but lower proportions in layer VI, (adjusted. *p* = 0.03; Fig. 4A, D). In contrast, density differences were pronounced and lateralised. RHCT demonstrated significantly higher L-NeuN+ density than RLCT across layers II, IV and VI (Fig 4G, I, J, adj. *p*=0.02 to <0.0001), with no corresponding differences in left TLE. RHCT also showed higher densities than LHCT in layers II, III and VI (Fig. 4G, H,J, adj. *p*=0.04 to 0.002). Two-way ANOVA confirmed a global increase in L-NeuN+ density across cortical layers in RHCT compared with all other groups (Fig. 4F). No left-right differences were observed in LCT groups.

**Figure 4.**
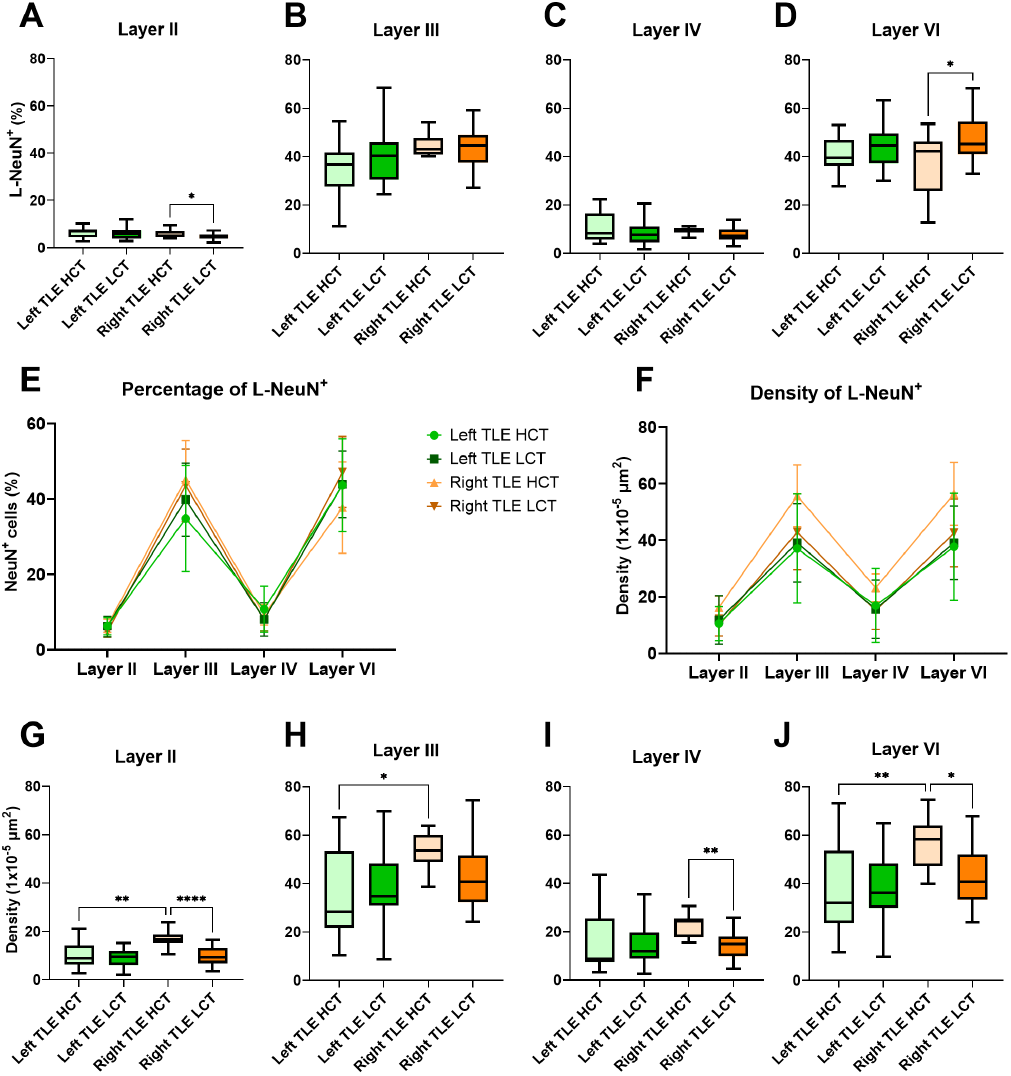
Morphometrics of distribution of Large NeuN^+^ cells between CT Groups in TLE-HS Type I cases. **A-D** shows one-way ANOVA analysis of Percentage L-NeuN^+^ between CT groups for each cortical layer, where comparisons are made between groups that had resection surgery on the same side and between same CT category, i.e. Left TLE HCT vs Right TLE HCT. Significant difference noted in Layer II and Layer VI for right sided cases with lower proportion of L-NeuN^+^ in Layer II and higher in Layer VI in LCT. (**E**) and (**F**) shows 2-way ANOVA analysis of repeat measures, accounting for multiple cortical layers measured within the same patient across CT groups. **(E)** depicts the Percentage L-NeuN^+^ and (**F**) Density L-NeuN^+^; significantly higher densities observed in Right HCT across layers (p = 0.01). **G-J** show one-way ANOVA of Density L-NeuN^+^ between groups, for each cortical layer, highlighting more consistent differences between left and right side in the HCT cases. All tests were corrected for multiple comparisons, significance is shown with asterisk, where * p<0.05, ** p<0.01, *** p<0.001 and **** p<0.0001. HCT = high cortical thinning; LCT = low cortical thinning; CT = cortical thinning.

#### S-NeuN^+^ distribution

Proportions of S-NeuN^+^ neurones were largely similar across groups, except for a higher percentage in RHCT compared with LHCT in layer III (adjusted *p* = 0.01; Fig. 5B). Density differences again showed strong lateralisation, with RHCT exhibiting higher S-NeuN^+^ density than RLCT across all layers reaching significance in layer II (adj. *p* = 0.002, Fig. 5G), IV (adj. *p* = 0.003, Fig. 5I) and VI (adj. *p* = 0.005, Fig. 5J). RHCT also showed higher densities than LHCT in layers II (adjusted *p* = 0.002; Fig. 5+G) and IV (adjusted *p* = 0.003; Fig. 5I). Again, there were no left-right differences in LCT groups.

**Figure 5.**
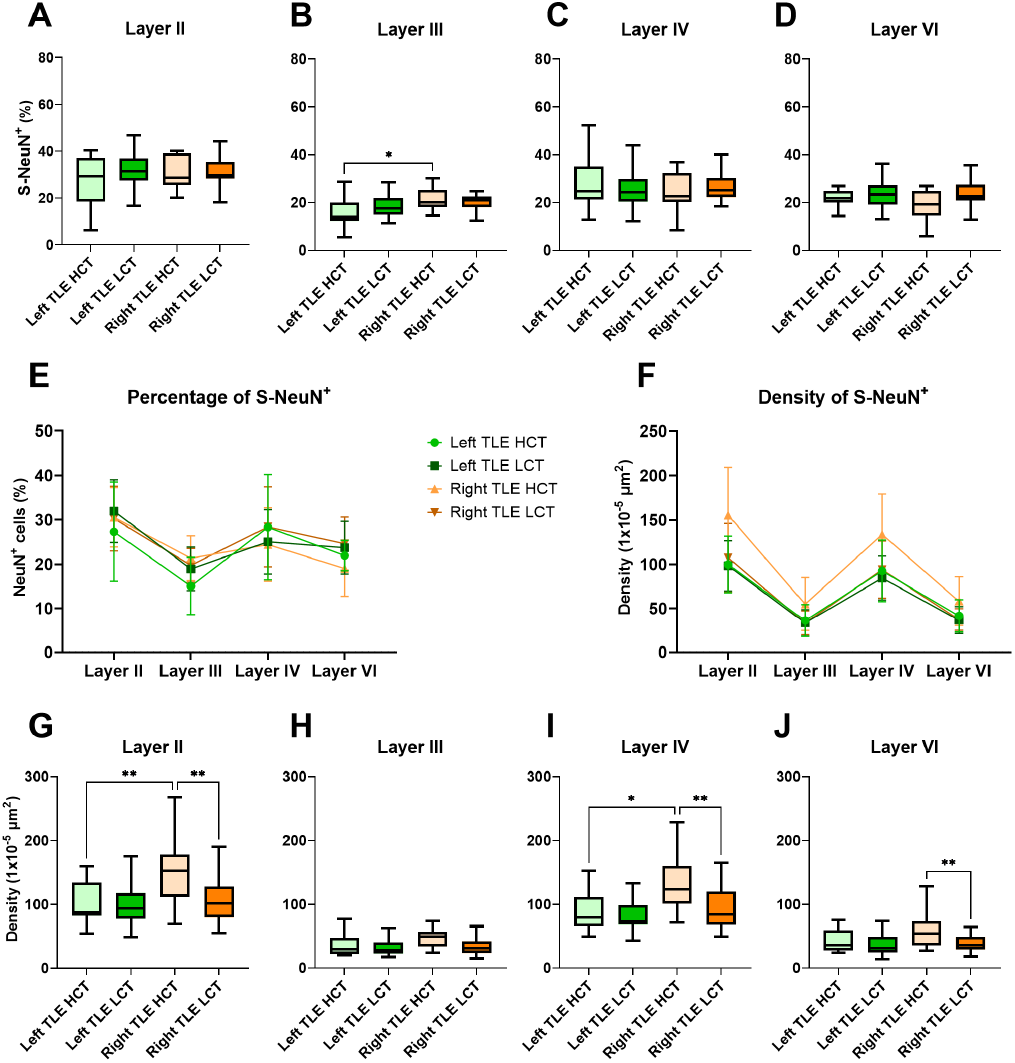
Morphometrics of distribution of Small NeuN+ cells between CT Groups in TLE-HS Type I cases. A-D shows one-way ANOVA analysis of Percentage S-NeuN+ between CT groups for each cortical layer, where comparisons are made between groups that had resection surgery on the same side and between same CT category, i.e. Left TLE HCT vs Right TLE HCT. (E) and (F) shows 2-way ANOVA analysis of repeat measures, accounting for multiple cortical layers measured within the same patient across CT groups. (E) depicts the Percentage S-NeuN+ and (F) Density S-NeuN+, significantly higher densities observed in Right HCT across layers (p < 0.001). G-J show one-way ANOVA of Density S-NeuN+ between groups, for each cortical layer, highlighting more consistent differences between left and right side in the HCT cases in Layers II (adj. p = 0.0003) and IV (adj. p = 0.01). All tests were corrected for multiple comparisons, significance is shown with asterisk, where * p<0.05 and ** p<0.01. HCT = high cortical thinning; LCT = low cortical thinning; CT = cortical thinning.

#### Sulcal-Gyral variations

Region-specific analyses across sulcal, crown and intermediate ROI revealed similar patterns in right TLE, although not uniformly across all regions (Fig. 6). Reduced layer VI thickness in RHCT compared with RLCT was confined only in the gyral crown (adj. *p*=0.006; Supplementary Fig. S9L), whereas increased L-NeuN^+^ density in RHCT was observed across superficial layers in all ROIs (Supplemental Figures S11). No significant differences were detected between LHCT and LLCT in any ROIs. Comparisons across ROIs within groups demonstrated greater sulcal variability in LCT than HCT cases bilaterally, suggesting reduced anatomical differentiation in HCT (Supplemental Fig. S12).

**Figure 6.**
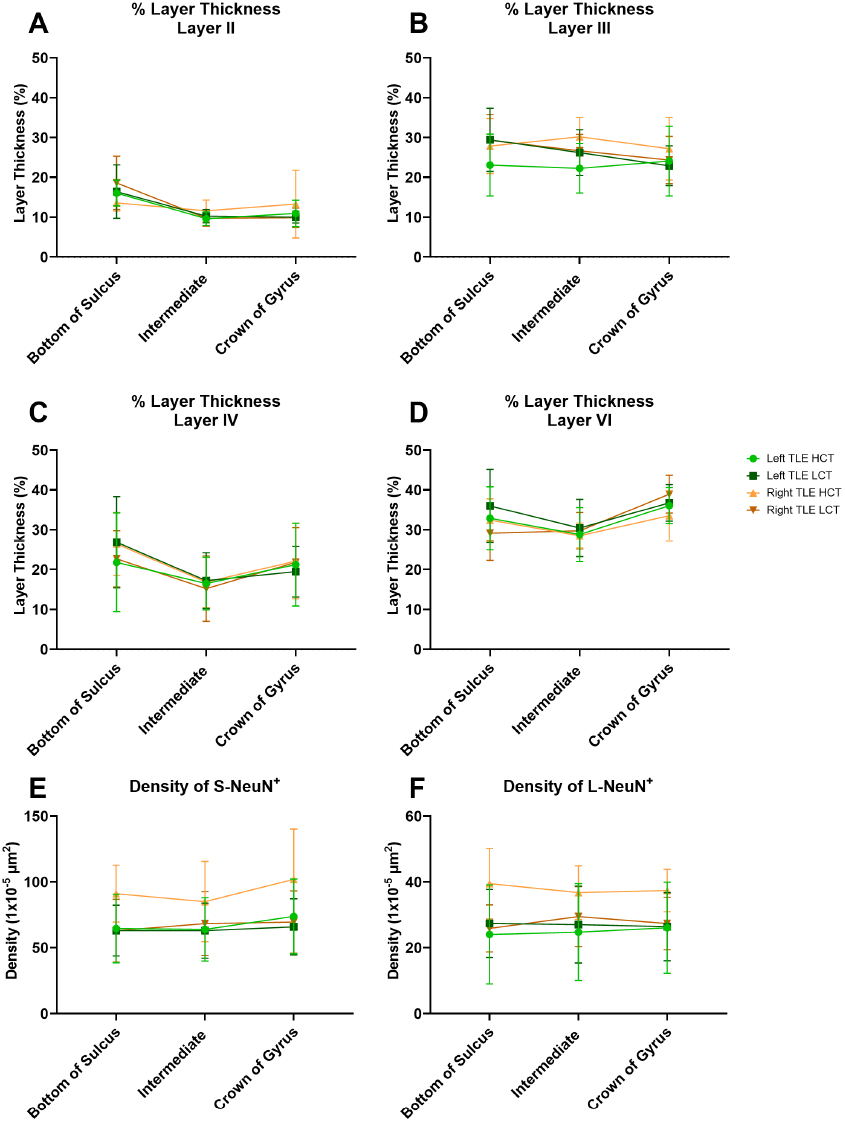
Comparison of dependent variables between ROIs. A-D shows repeat-measures 2-way ANOVA analysis of repeat measures of Layer Thickness for cortical layers: II (A), III (B), IV (C) and VI (D); (E) shows average density across all layers for S-NeuN+ and (F) density of L-NeuN+. A two-way ANOVA revealed a significant interaction between ROI and CT group on Layer Thickness for Layer II (p= 0.027; A), Layer III (p= 0.028; B), and Layer VI (p= 0.025; D). In (A), Layer II of RLCT had significantly higher layer thickness to RHCT at the bottom of sulcus (adj. p= 0.0295). For all CT groups, except RHCT, bottom of sulcus significantly differed from intermediate (LHCT p<0.001, LLCT p<0.001, RLCT p<0.001) and crown of gyrus (LHCT p<0.001, LLCT p<0.001, RLCT p<0.001). In (B), Layer III in intermediate region was significantly higher in RHCT than LHCT (p=0.04) and for both LCT groups, crown of gyrus significantly differed from bottom of sulcus (LLCT p=0.001, RLCT p=0.02) and intermediate (LLCT p=0.03). In C, Layer IV thickness remained similar for all groups between ROIs, however, within groups, bottom of sulcus significantly differed from intermediate (p=0.001) and crown of gyrus (p=0.007) in LLCT and intermediate differed significantly from bottom of sulcus (p<0.001) and crown of gyrus (p=0.008) in RLCT. All tests were corrected for multiple comparisons. In (D), Layer VI of LLCT had significantly higher layer thickness to RLCT at the bottom of sulcus (adj. p= 0.032). Intermediate region significantly differed from crown of gyrus in LLCT (p= 0.002), RHCT (p=0.02) and RLCT (p<0.001). In RLCT, bottom of sulcus also differed from crown of gyrus (p<0.001). All tests were corrected for multiple comparison. In summary, a two-way ANOVA of layer thickness revealed significant sulcal-gyral variations in the LCT but not HCT groups across regions in both hemispheres, implying diminished anatomical variations in layer thickness with cortical thinning. To view more details of this analysis, please see Supplementary Fig. S9. (E) two-way ANOVA revealed RHCT to have higher density of S-NeuN+ (not sig.), however, the main effect of both CT groups (adj. p = 0.0014) and ROI (adj. p < 0.0001) was significant. The density of S-NeuN+ in intermediate region was significantly lower than in crown of gyrus (adj. p = 0.05) in LHCT group and bottom of sulcus was significantly lower than crown of gyrus in LLCT (adj. p = 0.02). Intermediate region was significantly higher than both bottom of sulcus (adj. p = 0.008) and crown of gyrus (adj. p = 0.006) in RLCT. (F) showed that there is an overall effect of CT group on the density of L-NeuN+ (p=0.012). However, RHCT showed to have significantly higher density of L-NeuN+ in crown of gyrus over RLCT (p=0.005).

#### Regression Analysis: pathology-clinical measures

Regression analyses identified duration of epilepsy as a confounder due to its correlation with age at surgery and age at seizure onset (Supplementary Figs S13 and S14). After adjustment, layer thickness and neuronal proportions showed minimal associations with clinical variables (Supplementary Fig. S14B). However, higher densities of S-NeuN^+^ and L-NeuN^+^ neurones were positive predictors of MRI-defined cortical thickness in left TLE but negative predictors across all layers in right TLE. Age at surgery and age at onset also showed opposite associations with S-NeuN^+^ density between hemispheres. In left TLE only, poorer postoperative outcome was associated with higher L-NeuN^+^, across all layers (Supplementary Fig. S14). Focal to bilateral tonic-clonic seizures (FBTCS) was positively associated with higher L-NeuN^+^ in Layer III, when compared to focal impaired awareness seizures (FIAS) (*p=* 0.025; Supplementary Table S5). There was also a positive association of FBTCS type seizures with higher S-NeuN^+^ in layers II (*p=*0.029), IV (*p=*0.027) and VI (*p=*0.04). Conversely, there was a negative association between focal aware seizures (FAS) in relation to FIAS, for density of L-NeuN^+^ in layers II (*p=*0.02), III (*p<*0.001), IV (*p=*0.01) and VI (*p<*0.001); and for density of S-NeuN^+^ in layers II (*p<*0.001), III (*p=*0.05) and IV (*p=*0.003).

#### Automated vs Manual Annotation of Layers

Comparison of automated and semi-manual cortical layer delineation in LHCT cases showed excellent agreement on Bland-Altman analysis, with no systematic bias (Supplementary Fig. S15). The manual approach was more time-efficient (saving half the time) and is recommended for future studies.

## Discussion

We examined the histological substrates of MRI-defined cortical thinning in TLE using a semi-automated NeuN-based morphometric approach applied to a large surgical cohort. Across analyses, MRI-defined cortical thinning was not defined by overt laminar collapse or widespread neuronal loss. Rather, it was characterised by lateralised, layer-specific alterations in neuronal density and size distribution involving both superficial and deep cortical layers and most prominently in right-sided TLE with HS, with overall preservation of laminar architecture (Fig. 7).

**Figure 7.**
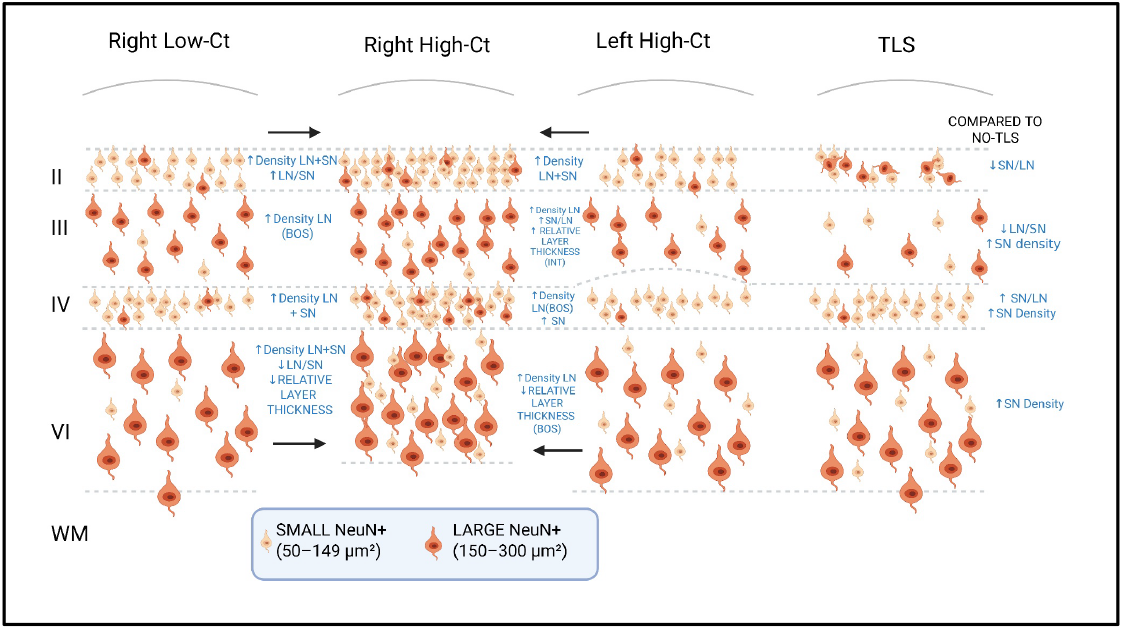
Diagrammatic representation of cytoarchitectural changes in relation to cortical thinning in study groups. The significant morphometric difference in cell-size distribution of NeuN+ cells on right high cortical thinning (HCT) cases, as categorised on MRI and with hippocampal sclerosis, is compared to left sided cases (HCT) and to right sides cases without cortical thinning (LCT). The key findings in temporal lobe sclerosis (TLS) are also shown, where morphometric findings identified cytoarchitectural changes in both superficial and deep cortical layers when compared to cases without TLS. Diagram made with Biorender.

Cortical thinning was also associated with attenuation of normal sulcal-gyral cytoarchitectural variations suggesting regional differentiation rather than uniform atrophy. Although our analyses were restricted to a single temporal gyrus and relied on two-dimensional, non-orthogonal sections, the findings support the notion that MRI-defined cortical thinning in TLE, unlike that observed in neurodegenerative disease, does not necessarily reflect irreversible neuronal loss.

### Cellular mechanisms underlying MRI-defined cortical thinning

Normal gradients of cortical thickness across the human brain are largely determined by the dimensions of layers III, V and VI and are closely linked to underlying function, connectivity and myeloarchitecture^39^, ranging approximately from 1.6-2.8 mm in the calcarine cortex to over 4.5 mm in the pre-central gyrus. The cortical volume is predominately composed of ‘neuropil’ (80-90%) made up of axons, dendrites, glia, vessels, extra cellular matrix and interstitial space which contributes substantially to thickness in addition to overall neuronal number^40^.

Taken together, our findings support a model in which MRI-defined cortical thinning reflects contraction of neuropil volume, manifesting as increased neuronal density, rather than neuronal depletion. This may be plausibly driven by changes in synaptic architecture, dendritic arborisation or myelin remodelling as cumulative effects of seizures^41,42^, although this requires further in-depth investigation.

The observed layer specific alterations in neuronal size distribution may represent adaptive or maladaptive responses to chronic network perturbation and altered microcircuit activity. Our findings are consistent with earlier stereological studies demonstrating preserved neuronal numbers with increased neuronal density and hypertrophy in the neocortex in TLE neocortex^25^, as well as more recent observations linking cortical thickness to neuronal size and packing rather than cell loss^31^. Emerging single-cell transcriptomic studies in refractory epilepsy, including TLE, further support this framework, identifying subsets of cortical pyramidal neurones with enlarged soma, mTOR pathway activation, altered dendritic architecture and stress- and senescence-related pathways^43^. Additional contributions from microglial activation^32^, particularly through synaptic pruning, and altered cortical myelin^26,40,44^ warrant further investigation. Furthermore, in this study only NeuN marker was used with cell-size classification to define laminar boundaries; future work incorporating cell-type specific markers will be necessary to determine whether particular neuronal subpopulations are selectively vulnerable.

Importantly, this neuropil- and cell size-based interpretation provides a biological explanation for the reported modulation or partial reversibility of MRI-defined cortical thinning following successful epilepsy surgery^17,18^, supporting the notion that at least a component of cortical thinning in TLE is dynamic rather than degenerative in nature.

### Methodological considerations in linking MRI cortical thickness to histology

Cortical thickness is defined as the Euclidean (shortest) distance between the pial and white matter boundaries^45^ and automated tools such as FreeSurfer and CAT12, demonstrate good reproducibility^46^. Identification of the grey-white boundary on MRI depends on the microstructural transition from high neuronal-rich cortex to axonal myelination, a contrast that may be reduced in TLE/HS, during development and with aging^47,48 49^. Normal regional variations in cortical thickness exist, reflecting underlying differences in cortical laminar cyto-architecture, myeloarchitectural and connectivity established during development^39,50^.

By focusing on the MTG at a defined coronal level we aimed to minimise anatomical variability and directly compare histological measures with contemporaneous pre-operative MRI-defined thickness measures. Previous histological studies of cortical thinning in epilepsy have been limited by small sample sizes and challenges inherent to two-dimensional, non-orthogonal tissue sections^25,31,51^. In our cohort, anatomical landmarks, such as radial vessels or apical dendrites on NeuN^+^ cells were inconsistently present for absolute cortical thickness estimation. To mitigate these limitations, we employed relative measures of laminar thickness and automated density-based layer delineation, which showed close agreement with manual measurements.

### Temporal lobe sclerosis and dissociation from cortical thinning

Temporal lobe sclerosis (TLS), a distinct cytoarchitectural alteration of the superficial cortex in a subset of TLE with HS^29^ provided an important internal control for our analyses. TLS cases were distributed across both high- and low cortical thinning groups and did not differ from no-TLS cases in MRI-defined cortical thickness or relative laminar thickness, confirming that TLS is not inevitably associated with cortical thinning, in keeping with earlier VBM studies^30^.

Nevertheless, TLS was characterised by predictable alterations in neuronal distribution in the superficial cortical layers, including reduced proportions of large neurones. Interestingly, we also observed significantly increased small neurone densities in deep cortical layers in TLS suggest this represents a more ‘pan-cortical’ developmental abnormality, warranting further in-depth study with neuronal lineage layer markers. Furthermore, given that TLS varies in its severity and spatial extent^26^, subtle or partial forms may be under-recognised on routine histopathological assessment. Automated cytoarchitectural profiling may prove useful for identifying TLS variants, particularly in light of reported associations with cognitive defects despite a similar electro-clinical phenotype to TLE/HS^52^.

### Lateralisation, systems-level organisation and clinical phenotype

A striking observation was the consistent lateralisation of cytoarchitectural alterations associated with MRI-defined cortical thinning. Right-sided TLE/HS showed more pronounced changes in neuronal density, altered laminar composition, reduced sulcal–gyral differentiation and selective thickness alterations involving layer VI, which is critical for descending cortico-thalamic output^53,54^ and these alterations may reflect corticothalamic signalling^9^. Notably, the thalamus has been reported to be reduced in volume in right TLE and negatively associated with epilepsy duration^6^. In contrast, left-sided TLE showed fewer structural alterations but stronger associations with clinical variables such as epilepsy duration and age at onset.

Normal hemispheric asymmetry of temporal lobe cortical thickness, with greater thickness on right, is well recognised^55^. Loss of this asymmetry has been linked to ageing and neurological disease^5^ and has been proposed as a marker of epileptogenic network vulnerability. Asymmetrical thinning in TLE have also attributed to developmental factors, including later maturation of the left hemisphere^56^. Notably, ENIGMA epilepsy consortium data indicate that inter-hemispheric asymmetry may dissociate from overall thinning patterns and is independent of disease duration and seizure onset^15^. Hemispheric functional specialisation provides an additional framework for interpreting these findings. The left temporal lobe, dominant for language in most individuals, is subject to sustained use-dependent activation across the lifespan. Cognitive trajectories in TLE reflect an interaction between early neurodevelopmental compromise and later network reorganisation rather than uniform progressive degeneration. The hemispheric dominance and functional reserve shape vulnerability patterns, with differential cognitive and structural consequences in left versus right TLE^57^. Within this framework, relative structural preservation of the left temporal cortex in some patients may reflect activity-dependent stabilisation of language networks or prioritisation of functionally critical circuitry. This interpretation is also consistent with the known ageing-related preferential decline of right-hemispheric functions, potentially preserving dominant language-related systems.

In our cohort, anticipated right–left asymmetry was absent in cases with hippocampal sclerosis but without MRI-defined cortical thinning, suggesting that pathological reorganisation may precede or occur independently of measurable atrophy. The distinct pathological-clinical associations between hemispheres further support the concept that left and right TLE represent partially distinct biological syndromes^6,36^, potentially reflecting different developmental and network trajectories. In left TLE, higher neuronal densities were negatively associated with epilepsy duration, consistent with MRI studies linking cortical thinning to seizure burden^12,13^, whereas, right TLE showed weaker or opposing associations, reinforcing the notion of lateralised pathophysiological mechanisms.

### Limitations

This study has several limitations. Histological analyses were restricted to a single temporal gyrus and relied on two-dimension, non-orthogonal sections, necessitating the use of relative rather than absolute measures. Pre-operative neuropsychological data were not available for all cases and will be addressed in a future study. Recent MRI studies using event-based and subtype modelling approaches demonstrate heterogenous disease trajectories in TLE^4,45^, which were not explicitly modelled here and may contribute to variability in effect size. In addition, no healthy control group was available for direct comparison, as post-mortem tissue does not stain consistently well with NeuN for our image analysis method and bilateral comparisons within individual surgical cases were not possible.

## Conclusion

MRI-defined cortical thinning in TLE is not a simple marker of neuronal loss or laminar atrophy. Rather, cortical thinning reflects lateralised alterations in neuronal density, size distribution and cortical organisation, particularly in right-sided TLE, alongside reduced sulcal– gyral differentiation and some evidence of more selective involvement of layer VI. These changes are more consistent with structural reorganisation of cortical microcircuits and network-level pathology than with irreversible neurodegeneration. Recognising the biological complexity underlying MRI-defined cortical thinning has important implications for its interpretation as a biomarker for understanding disease mechanisms and importantly its potential reversibility in temporal lobe epilepsy.

## Data availability

The data and QuPath script used in this study are available from the corresponding author, upon reasonable request.

## Acknowledgements

We would like to acknowledge support from Pan Shengning from UCL Statistics Group.

## References

1. Dickerson BC, Bakkour A, Salat DH, et al. The Cortical Signature of Alzheimer’s Disease: Regionally Specific Cortical Thinning Relates to Symptom Severity in Very Mild to Mild AD Dementia and is Detectable in Asymptomatic Amyloid-Positive Individuals. Cerebral Cortex. 2009;19(3):497–510. doi:10.1093/cercor/bhn113

2. Pacheco J, Goh JO, Kraut MA, Ferrucci L, Resnick SM. Greater cortical thinning in normal older adults predicts later cognitive impairment. Neurobiology of Aging. 2015;36(2):903–908. doi:10.1016/j.neurobiolaging.2014.08.031

3. Wong TY, Radua J, Pomarol-Clotet E, et al. An overlapping pattern of cerebral cortical thinning is associated with both positive symptoms and aggression in schizophrenia via the ENIGMA consortium. Psychol Med. 2020;50(12):2034–2045. doi:10.1017/S0033291719002149

4. Xiao F, Caciagli L, Wandschneider B, et al. Identification of different MRI atrophy progression trajectories in epilepsy by subtype and stage inference. Brain. 2023;146(11):4702–4716. doi:10.1093/BRAIN/AWAD284

5. Roe JM, Vidal-Piñeiro D, Sørensen Ø, et al. Asymmetric thinning of the cerebral cortex across the adult lifespan is accelerated in Alzheimer’s disease. Nat Commun. 2021;12(1):721. doi:10.1038/s41467-021-21057-y

6. Whelan CD, Altmann A, Botía JA, et al. Structural brain abnormalities in the common epilepsies assessed in a worldwide ENIGMA study. Brain. 2018;141(2):391–408. doi:10.1093/brain/awx341

7. Galovic M, Van Dooren VQH, Postma TS, et al. Progressive Cortical Thinning in Patients with Focal Epilepsy. JAMA Neurology. 2019;76(10):1230–1239. doi:10.1001/jamaneurol.2019.1708

8. Kaestner E, Reyes A, Chen A, et al. Atrophy and cognitive profiles in older adults with temporal lobe epilepsy are similar to mild cognitive impairment. Brain. 2021;144(1):236–250. doi:10.1093/brain/awaa397

9. Keller SS, Richardson MP, Schoene-Bake JC, et al. Thalamotemporal alteration and postoperative seizures in temporal lobe epilepsy. Annals of Neurology. 2015;77(5):760–774. doi:10.1002/ana.24376

10. McDonald CR, Hagler Jr DJ, Ahmadi ME, et al. Regional neocortical thinning in mesial temporal lobe epilepsy. Epilepsia. 2008;49(5):794–803. doi:10.1111/j.1528-1167.2008.01539.x

11. Mueller SG, Laxer KD, Barakos J, Cheong I, Garcia P, Weiner MW. Widespread neocortical abnormalities in temporal lobe epilepsy with and without mesial sclerosis. NeuroImage. 2009;46(2):353–359. doi:10.1016/j.neuroimage.2009.02.020

12. Lin JJ, Salamon N, Lee AD, et al. Reduced Neocortical Thickness and Complexity Mapped in Mesial Temporal Lobe Epilepsy with Hippocampal Sclerosis. Cereb Cortex. 2007;17(9):2007–2018. doi:10.1093/cercor/bhl109

13. Bernhardt BC, Worsley KJ, Kim H, Evans AC, Bernasconi A, Bernasconi N. Longitudinal and cross-sectional analysis of atrophy in pharmacoresistant temporal lobe epilepsy. Neurology. 2009;72(20):1747–1754. doi:10.1212/01.wnl.0000345969.57574.f5

14. Larivière S, Rodríguez-Cruces R, Royer J, et al. Network-based atrophy modeling in the common epilepsies: A worldwide ENIGMA study. Sci Adv. 2020;6(47):eabc6457. doi:10.1126/sciadv.abc6457

15. Park B yong, Larivière S, Rodríguez-Cruces R, et al. Topographic divergence of atypical cortical asymmetry and atrophy patterns in temporal lobe epilepsy. Brain. 2022;145(4):1285–1298. doi:10.1093/brain/awab417

16. Chen J, Ngo A, Rodriguez-Cruces R, et al. Changes in Gray Matter Morphology and White Matter Microstructure Across the Adult Lifespan in People With Temporal Lobe Epilepsy. Neurology. 2025;105(6):e213688. doi:10.1212/WNL.0000000000213688

17. Galovic M, de Tisi J, McEvoy A, et al. Resective surgery prevents progressive cortical thinning in temporal lobe epilepsy. Brain. 2020;143(11):3172–3175. doi:10.1093/BRAIN/AWAA330

18. Li W, Jiang Y, Qin Y, et al. Cortical remodeling before and after successful temporal lobe epilepsy surgery. Acta Neuro Scandinavica. 2022;146(2):144–151. doi:10.1111/ane.13631

19. Keller SS, Roberts N. Voxel-based morphometry of temporal lobe epilepsy: An introduction and review of the literature. Epilepsia. 2008;49(5):741–757. doi:10.1111/j.1528-1167.2007.01485.x

20. Ogren JA, Tripathi R, Macey PM, et al. Regional cortical thickness changes accompanying generalized tonic-clonic seizures. NeuroImage: Clinical. 2018;20:205–215. doi:10.1016/j.nicl.2018.07.015

21. Sartorius A, Demirakca T, Böhringer A, et al. Electroconvulsive therapy increases temporal gray matter volume and cortical thickness. European Neuropsychopharmacology. 2016;26(3):506–517. doi:10.1016/j.euroneuro.2015.12.036

22. Won J, Gogniat M, Smith JC. Age and Sex-Related Differences in Neuroprotective Effects of Cardiovascular Endurance on Cortical Thickness and Brain Volume in Adults Across Age. Brain and Behavior. 2025;15(1):e70231. doi:10.1002/brb3.70231

23. Liu J, Reeves C, Jacques T, et al. Nestin-expressing cell types in the temporal lobe and hippocampus: Morphology, differentiation, and proliferative capacity. GLIA. 2018;66(1):62–77. doi:10.1002/glia.23211

24. Liu J, Binding L, Puntambekar I, et al. Microangiopathy in temporal lobe epilepsy with diffusion MRI alterations and cognitive decline. Acta Neuropathol. 2024;148(1):49. doi:10.1007/s00401-024-02809-8

25. Bothwell S, Meredith GE, Phillips J, et al. Neuronal Hypertrophy in the Neocortex of Patients with Temporal Lobe Epilepsy. J Neurosci. 2001;21(13):4789–4800. doi:10.1523/JNEUROSCI.21-13-04789.2001

26. Thom M, Eriksson S, Martinian L, et al. Temporal lobe sclerosis associated with hippocampal sclerosis in temporal lobe epilepsy: Neuropathological features. Journal of Neuropathology and Experimental Neurology. 2009;68(8):928–938. doi:10.1097/NEN.0b013e3181b05d67

27. Eriksson SH, Free SL, Thom M, et al. Correlation of quantitative MRI and neuropathology in epilepsy surgical resection specimens—T2 correlates with neuronal tissue in gray matter. NeuroImage. 2007;37(1):48–55. doi:10.1016/j.neuroimage.2007.04.051

28. Lorio S, Kherif F, Ruef A, et al. Neurobiological origin of spurious brain morphological changes: A quantitative MRI study. Human Brain Mapping. 2016;37(5):1801–1815. doi:10.1002/hbm.23137

29. Najm I, Lal D, Alonso Vanegas M, et al. The ILAE consensus classification of focal cortical dysplasia: An update proposed by an ad hoc task force of the ILAE diagnostic methods commission. Epilepsia. 2022;63(8):1899–1919. doi:10.1111/epi.17301

30. Eriksson SH, Thom M, Symms MR, et al. Cortical neuronal loss and hippocampal sclerosis are not detected by voxel-based morphometry in individual epilepsy surgery patients. Human Brain Mapping. 2009;30(10):3351–3360. doi:10.1002/hbm.20757

31. Heyer DB, Wilbers R, Galakhova AA, et al. Verbal and General IQ Associate with Supragranular Layer Thickness and Cell Properties of the Left Temporal Cortex. Cerebral Cortex. 2022;32(11):2343–2357. doi:10.1093/cercor/bhab330

32. Altmann A, Ryten M, Di Nunzio M, et al. A systems-level analysis highlights microglial activation as a modifying factor in common epilepsies. Neuropathology Appl Neurobio. 2021;48(1):e12758. doi:10.1111/nan.12758

33. Najm IM, Sarnat HB, Blümcke I. Review: The international consensus classification of Focal Cortical Dysplasia – a critical update 2018. Neuropathology Appl Neurobio. 2018;44(1):18–31. doi:10.1111/nan.12462

34. Gaser C, Dahnke R, Thompson PM, Kurth F, Luders E, the Alzheimer’s Disease Neuroimaging Initiative. CAT: a computational anatomy toolbox for the analysis of structural MRI data. Gigascience. 2024;13:giae049. doi:10.1093/gigascience/giae049

35. Zhang C, Hu W, Wu Y, Li G, Yang C, Wu T. Altered Directed-Connectivity Network in Temporal Lobe Epilepsy: A MEG Study. Sensors. 2025;25(5):1356. doi:10.3390/s25051356

36. Blumcke I, Spreafico R, Haaker G, et al. Histopathological Findings in Brain Tissue Obtained during Epilepsy Surgery. N Engl J Med. 2017;377(17):1648–1656. doi:10.1056/NEJMoa1703784

37. Eriksson SH, Free SL, Thom M, Harkness W, Sisodiya SM, Duncan JS. Reliable Registration of Preoperative MRI with Histopathology after Temporal Lobe Resections. Epilepsia. 2005;46(10):1646–1653. doi:10.1111/j.1528-1167.2005.00257.x

38. Bankhead P, Loughrey MB, Fernández JA, et al. QuPath: Open source software for digital pathology image analysis. Sci Rep. 2017;7(1):16878. doi:10.1038/s41598-017-17204-5

39. Wagstyl K, Ronan L, Goodyer IM, Fletcher PC. Cortical thickness gradients in structural hierarchies. NeuroImage. 2015;111:241–250. doi:10.1016/j.neuroimage.2015.02.036

40. Wagstyl K, Larocque S, Cucurull G, et al. BigBrain 3D atlas of cortical layers: Cortical and laminar thickness gradients diverge in sensory and motor cortices. Kennedy H, ed. PLoS Biol. 2020;18(4):e3000678. doi:10.1371/journal.pbio.3000678

41. Li H, Xie Z, Tian Y, et al. Genome-wide consensus transcriptional signatures identify synaptic pruning linking Alzheimer’s disease and epilepsy. Mol Psychiatry. Published online October 26, 2025. doi:10.1038/s41380-025-03318-0

42. Kumari S, Brewster AL. Exploring Dendritic and Spine Structural Profiles in Epilepsy: Insights From Human Studies and Experimental Animal Models. Epilepsy Curr. 2024;24(1):40–46. doi:10.1177/15357597231218603

43. Ge Q, Yang J, Huang F, et al. Multimodal single-cell analyses reveal molecular markers of neuronal senescence in human drug-resistant epilepsy. Journal of Clinical Investigation. 2025;135(5):e188942. doi:10.1172/JCI188942

44. Bernhardt BC, Fadaie F, Vos De Wael R, et al. Preferential susceptibility of limbic cortices to microstructural damage in temporal lobe epilepsy: A quantitative T1 mapping study. NeuroImage. 2018;182:294–303. doi:10.1016/j.neuroimage.2017.06.002

45. Lee HM, Fadaie F, Gill RS, et al. MRI-Derived Modeling of Disease Progression Patterns in Patients With Temporal Lobe Epilepsy. Neurology. 2024;103(3). doi:10.1212/WNL.0000000000209524

46. Dias MDFM, Carvalho P, Castelo-Branco M, Valente Duarte J. Cortical thickness in brain imaging studies using FreeSurfer and CAT12: A matter of reproducibility. Neuroimage: Reports. 2022;2(4):100137. doi:10.1016/j.ynirp.2022.100137

47. Garbelli R, Milesi G, Medici V, et al. Blurring in patients with temporal lobe epilepsy: clinical, high-field imaging and ultrastructural study. Brain. 2012;135(8):2337–2349. doi:10.1093/brain/aws149

48. Rodríguez-Cruces R, Concha L. White matter in temporal lobe epilepsy: clinico-pathological correlates of water diffusion abnormalities. Quantitative Imaging in Medicine and Surgery. 2015;5(2):26478–26278. doi:10.3978/j.issn.2223-4292.2015.02.06

49. Song JY, Fleysher R, Ye K, et al. Characterizing the microstructural transition at the gray matter-white matter interface: Implementation and demonstration of age-associated differences. NeuroImage. 2025;306:121019. doi:10.1016/j.neuroimage.2025.121019

50. Huntenburg JM, Bazin PL, Margulies DS. Large-Scale Gradients in Human Cortical Organization. Trends in Cognitive Sciences. 2018;22(1):21–31. doi:10.1016/j.tics.2017.11.002

51. Cardinale F, Chinnici G, Bramerio M, et al. Validation of FreeSurfer-Estimated Brain Cortical Thickness: Comparison with Histologic Measurements. Neuroinform. 2014;12(4):535–542. doi:10.1007/s12021-014-9229-2

52. Cossu M, d’Orio P, Barba C, et al. Focal Cortical Dysplasia IIIa in Hippocampal Sclerosis-Associated Epilepsy: Anatomo-Electro-Clinical Profile and Surgical Results From a Multicentric Retrospective Study. Neurosurg. 2021;88(2):384–393. doi:10.1093/neuros/nyaa369

53. Jones EG. Thalamic circuitry and thalamocortical synchrony. Adams P, Guillery RW, Sherman SM, Sillito AM, eds. Phil Trans R Soc Lond B. 2002;357(1428):1659–1673. doi:10.1098/rstb.2002.1168

54. Bomben VC, Aiba I, Qian J, Mark MD, Herlitze S, Noebels JL. Isolated P/Q Calcium Channel Deletion in Layer VI Corticothalamic Neurons Generates Absence Epilepsy. J Neurosci. 2016;36(2):405–418. doi:10.1523/JNEUROSCI.2555-15.2016

55. Kong XZ, Mathias SR, Guadalupe T, et al. Mapping cortical brain asymmetry in 17,141 healthy individuals worldwide via the ENIGMA Consortium. Proc Natl Acad Sci USA. 2018;115(22). doi:10.1073/pnas.1718418115

56. Kemmotsu N, Girard HM, Bernhardt BC, et al. MRI analysis in temporal lobe epilepsy: Cortical thinning and white matter disruptions are related to side of seizure onset: Structural Compromise in TLE. Epilepsia. 2011;52(12):2257–2266. doi:10.1111/j.1528-1167.2011.03278.x

57. Helmstaedter C, Elger CE. Chronic temporal lobe epilepsy: a neurodevelopmental or progressively dementing disease? Brain. 2009;132(10):2822–2830. doi:10.1093/brain/awp182

